# *In situ* structural determination of cyanobacterial phycobilisome-PSII supercomplex by STAgSPA strategy

**DOI:** 10.1101/2023.12.17.572042

**Authors:** Xing Zhang, Yanan Xiao, Xin You, Shan Sun, Sen-Fang Sui

**Affiliations:** Ministry of Education Key Laboratory of Protein Sciences, Tsinghua-Peking Joint Center for Life Sciences, Beijing Frontier Research Center for Biological Structures, Beijing Advanced Innovation Center for Structural Biology, School of Life Sciences, Tsinghua University, Beijing 100084, China; School of Life Sciences, Southern University of Science and Technology, Shenzhen 518055, Guangdong, China; State Key Laboratory of Membrane Biology, Beijing Frontier Research Center for Biological Structures, Beijing Advanced Innovation Center for Structural Biology, School of Life Sciences, Tsinghua University, Beijing 100084, China

## Abstract

Photosynthesis converting solar energy to chemical energy is one of the most important chemical reactions on earth^1^. In cyanobacteria, light energy is captured by antenna system phycobilisomes (PBSs) and transferred to photosynthetic reaction centers of photosystem II (PSII) and photosystem I (PSI)^2^. While most of the protein complexes involved in photosynthesis have been characterized by *in vitro* structural analyses, how these protein complexes function together *in vivo* is not well understood. Here we developed an *in situ* structural analysis strategy “STAgSPA” to successfully solve the *in situ* structure of PBS-PSII supercomplex from the cyanobacteria *Spirulina platensis* FACHB-439 at resolution of ∼3.5Å. The structure reveals the unprecedented coupling details among adjacent PBSs and PSII dimers, and the collaborative energy transfer mechanism mediated by multiple “super-PBS” in cyanobacteria. Our results not only provide the insights for understanding the diversity of photosynthesis-related systems between prokaryotic cyanobacteria and eukaryotic red algae, but also a valuable methodological demonstration for *in situ* high-resolution structural analysis in cellular or tissue samples.

## Introduction

In oxygenic photosynthetic organisms, the light-driven reactions depend on the cooperation of different multiprotein complexes that attached on or embedded in thylakoid membranes, such as antenna systems, photosystems II/I (PSII/PSI) and cytochrome *b_6_f*^3,4^. In cyanobacteria, sunlight is harvested by the soluble light-harvesting apparatus phycobilisomes (PBSs), and the energy is finally transferred to the reaction center of PSII/PSI to induce the photo-induced electron transport^2,5^. The structures of isolated PBS, PSII and PSI have been resolved individually by cryo-electron microscopy (cryo-EM) single-particle analysis (SPA) at near-atomic resolution^6–9^, providing the structural basis for energy transfer, electron transfer, and photoprotection within the isolated complexes. However, it is hard to obtain the intact assemblies of these protein machineries by SPA due to their diverse protein properties, which may result in the dissociation of some loosely associated supercomplexes during purification. Although spectroscopic, the cross-linking coupled with mass spectrometry (CXMS) analysis and some low-resolution negative-staining EM structures have identified several interlinks and coarse binding pattern between PBS and PSII/PSI in cyanobacteria^10–12^, the precise interaction and energy transfer pathways at native state still remain unclear.

Cryo-electron tomography (Cryo-ET) coupled with subtomogram averaging (STA) emerges as a potent methodology for addressing target particles in cellular environment, which has been applied to visualize arrangement of PBSs, PSII or ATPase in thylakoid membranes^13,14^. However, achieving sub-nanometer resolution through the standard procedure of cryo-ET-STA is still difficult due to a series of limiting factors, including limited amount of collected data, low signal-to-noise ratio, inaccurate tilt-series alignment, etc. Thus, the utilization of the typical pipeline of cryo-ET-STA only provides the cellular-scale observation of the binding of the PBS and PSII dimer with stoichiometry 1:1 from cyanobacteria *Synechocystis* without structural detail^14^. Recently, single-particle analysis was also applied to *in situ* structural analysis through a high-resolution template matching procedure (isSPA), which emerged as an alternative approach and has successfully solved several *in situ* structures at near-atomic resolution^15–19^. However, a prerequisite that a high-resolution template is needed for particle detection significantly limits the application prospects of this method. To overcome these challenges, we have developed a new *in situ* structural analysis strategy that combined cryo-ET-STA and SPA techniques to determine the *in situ* structures in the cellular context, termed “<u>s</u>ubtomogram <u>a</u>veraging <u>g</u>uided <u>s</u>ingle-<u>p</u>article <u>a</u>nalysis (STAgSPA)”. We have successfully applied this strategy to solve the *in situ* structure of the PBS-PSII-PSI-LHC megacomplexes from red alga *Porphyridium purpureum* at an overall resolution of 3.3 Å^20^. The *in situ* high-resolution structure reveals the precise association and bilin distribution among PBS, PSII and PSI of red algae. Since the native photosynthetic apparatuses are compositionally and structurally diverse, *in situ* structural analysis in other species are essential for the comprehensive understanding of their working mechanisms.

Herein, we further employed cryo-focused ion beam (cryo-FIB) milling and the STAgSPA to address the *in situ* structure of the cyanobacterial PBS-PSII supercomplex from *Spirulina platensis*, a more challenging sample due to the smaller size and more flexible property of cyanobacterial PBS compared to the red algal PBS. We finally determined the structure at resolution of ∼3.5 Å, demonstrating the powerful capability of the STAgSPA strategy to deal with complicated sample and to push the *in situ* structural determination to near-atomic level. The structure reveals unprecedented details concerning the specific interactions between PBS and PSII in cyanobacteria, and a huge number of pigment networks that contribute to energy transfer. Moreover, we conducted comparative analysis between the prokaryotic cyanobacterial PBS-PSII supercomplex and the eukaryotic red algal PBS-PSII module reported recently^20^, aiming to elucidate both their shared characteristics and distinctive features. This investigation provides valuable insights into the evolutionary perspective of light energy capture and transfer in photosynthetic organisms.

## Results and discussion

### STAgSPA, an *in situ* structural analysis strategy

As shown in figure 1, the STAgSPA workflow contains: (1) lamellae production by cryo-FIB milling, and the coarse screening by TEM in low magnifications is used to retain intact, no ice contamination and well-thickness lamellae (Fig. 1a); (2) data collection of the cryo-FIB lamellae is performed in two different ways. Some lamellae are used to collect the typical tilt series dataset, and others are used for the high-dose zero-tilt dataset collection (Fig. 1b); (3) for the tilt data set, tomogram reconstruction and STA are performed to generate an intermediate-resolution structure of the target, which will be used as a template in the next step (Fig. 1c); (4) using the isSPA method^15,16^, target particle detection is carried out for the high-dose zero-tilt images, yielding initial coordinates and Euler angles (Fig. 1d); (5) the target particles are further processed to constrained single-particle refinement and 3D classification to generate the final high-resolution reconstruction (Fig. 1d).

**Fig. 1.**
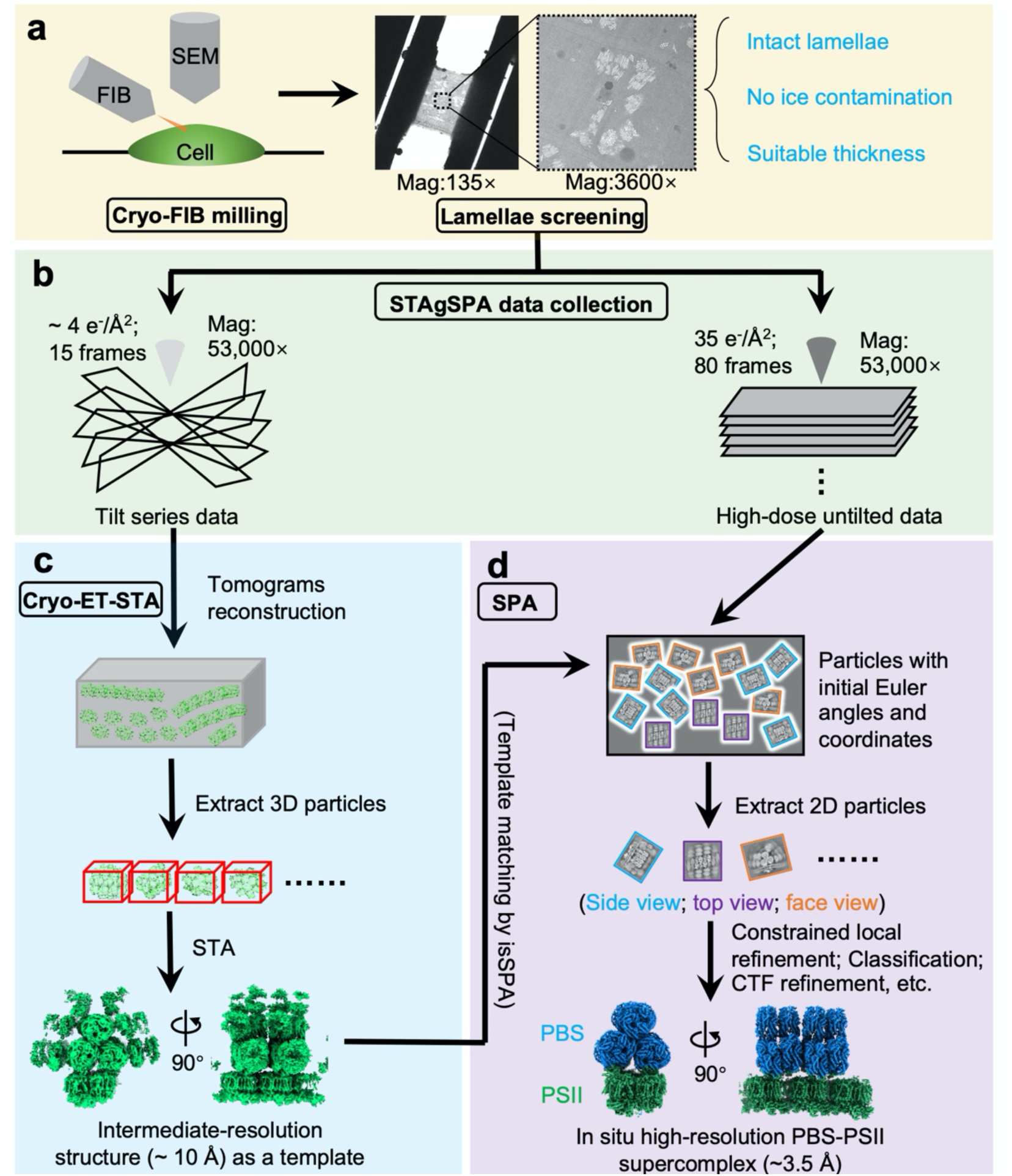
Overview of the STAgSPA workflow. **a**, Cell lamellae are produced by cryo-FIB milling. The coarse screening by TEM in low magnifications is used to retain intact, no ice contamination and well-thickness lamellae. **b**, STAgSPA data collection strategy. The coarse screened lamellae are divided into two parts, one for tilt series data collection (∼ 4 e^-^/Å^2^ per image with 15 frames) and the other one for high-dose untilted data collection (∼ 35 e^-^/Å^2^ per image with 80 frames). **c**, A conventional Cryo-ET-STA workflow containing tomogram reconstruction, particle picking and subtomogram averaging, is applied to tilt series data to yield an intermediate-resolution structure of the PBS-PSII supercomplex (∼ 10 Å) used as a template in the further SPA step. **d**, high-dose untilted data is used to generate the high-resolution structure by SPA workflow, including particle picking via template matching by isSPA, constrained local refinement and 3D classification, et.al. Using STAgSPA strategy, we solved the *in situ* structure of the PBS-PSII supercomplex from cyanobacteria *S. platensis* at the resolution of ∼ 3.5 Å.

Notably, our approach differs from the previously reported hybrid STA-SPA method, in which the high-dose image was first acquired at 0° followed by the typical tilt series data collection on the same piece of lamella, which is time-consuming^21,22^. In STAgSPA, the high-dose zero-tilt images are acquired separately from the tilt series and in the way same as the single particle data collection, which is very productive and highly efficient. Moreover, because of the limitation of total electron exposure dose for biological sample, increased electron dose for zero-tilt images in the hybrid STA-SPA method results in the decreased electron dose for tilt images, which leads to the low signal-to-noise ratio and thus the negative impacts on the cryo-ET-STA data processing. This problem is avoided in STAgSPA, as the high-dose single-particle images and the tomographic tilt images are collected from different lamellae with normal electron doses. Furthermore, our method addresses the reference selection problem effectively by incorporating intermediate-resolution STA outputs as templates within the isSPA particle detection^15,16^. STAgSPA also offers advantages over other options: (1) When using simulated EM density derived from homology PDB models, there may be mismatches between the template and the target protein. Such mismatches can affect detection efficiency and increase the risk of template bias; (2) If employing reported density maps from single-particle analysis conducted on different cryo-EM imaging systems, differences in protein conformation or errors in imaging parameters (such as pixel size) could also hinder the accurate detection.

In this research we have attempted to detect the target particles in the high-dose images using only a single cyanobacterial high-resolution PBS structure solved by cryo-EM SPA and a simulated density generated by PDB model in the isSPA method^7,23^, but the particle detection was failed. This may be because PBSs are arranged into arrays in cellular state, using individual PBS density may not fully harness the signals. We finally used the STAgSPA strategy, in which an intermediate-resolution three-layer PBS array from STA served as an optimal template to guide the particle picking, alignment and refinement during the single-particle data processing, to obtain the structure at the near-atomic resolution of ∼3.5 Å. Our results demonstrate that STAgSPA could effectively circumvent limitations of using cryo-ET-STA or isSPA alone^15,16^, and thus advance the *in situ* structural analysis into the high-resolution level.

### Overall structure of *in situ* cyanobacterial PBS-PSII supercomplex

The cyanobacterium *S. platensis* was cultured under continuous low orange light illumination at 25 °C with an intensity of ∼ 10 μmol of photons m^-2^ s^-1^ and the details of cryo-sample preparation, cryo-FIB milling, image collection and data processing by STAgSPA approach are described in the Methods. Briefly, cells in exponential growth phase were harvested and vitrified on cryo-EM grids after 20 min dark-adaption. Cryo-FIB was used to produce thin lamellas with ∼150 nm thickness for collecting typical tilt series and zero-tilt high-dose images. Subsequently, the tilt series were processed by cryo-ET-STA workflow to generate the template used in particle detection by isSPA method for the zero-tilt images. After multi-step focused SPA refinement, the structure of the PBS-PSII supercomplex was yielded (Fig. 1, Extended Data Fig. 1 and 2). The local resolutions of the subcomplexes are 3.6 Å and 3.5 Å for PBS and PSII dimer, respectively (Extended Data Fig. 2). The overall structure of cyanobacterial PBS-PSII supercomplex has dimensions of approximately 360 × 330 × 380 Å^3^ with a total mass of approximately 7.6 MDa (Fig. 2a). The structure shows that two face-to-face stacked PBSs (labeled as PBS-1 and PBS-2) anchor to three PSII dimers (PSII-d1, -d2 and -d3) with a 1:1 stoichiometry between PBS and PSII dimer, and repeat periodically along the PSII array of thylakoid membrane (Fig. 2a-c). Each PBS spans two PSII dimers with an angle of 6° between the hemi-discoidal plane of the PBS core and the central plane of the PSII dimer pair, and the PBS center is offset by nearly 4.5 nm relative to the PSII dimer pair center (Fig. 2c). This misalignment between the centers of PBS and PSII suggests that the two pairs of symmetric terminal emitters of each PBS, ApcD/ApcD’ and L_CM_/L_CM_’, contact with PSII dimers in different manners. Moreover, two adjacent PBS cores show parallel-displaced organization with a ∼5 nm shift along the PBS array (Fig. 2c). Notably, red algal PBS-PSII exhibits a larger rotation angle (∼14°) but no PBS core offset relative to the PSII dimer pair^20,24^, suggesting PBS-PSII supercomplex from the prokaryotic cyanobacteria may have distinctive strategies for association and energy transfer pathways between PBS and photosystems (described below).

**Fig. 2.**
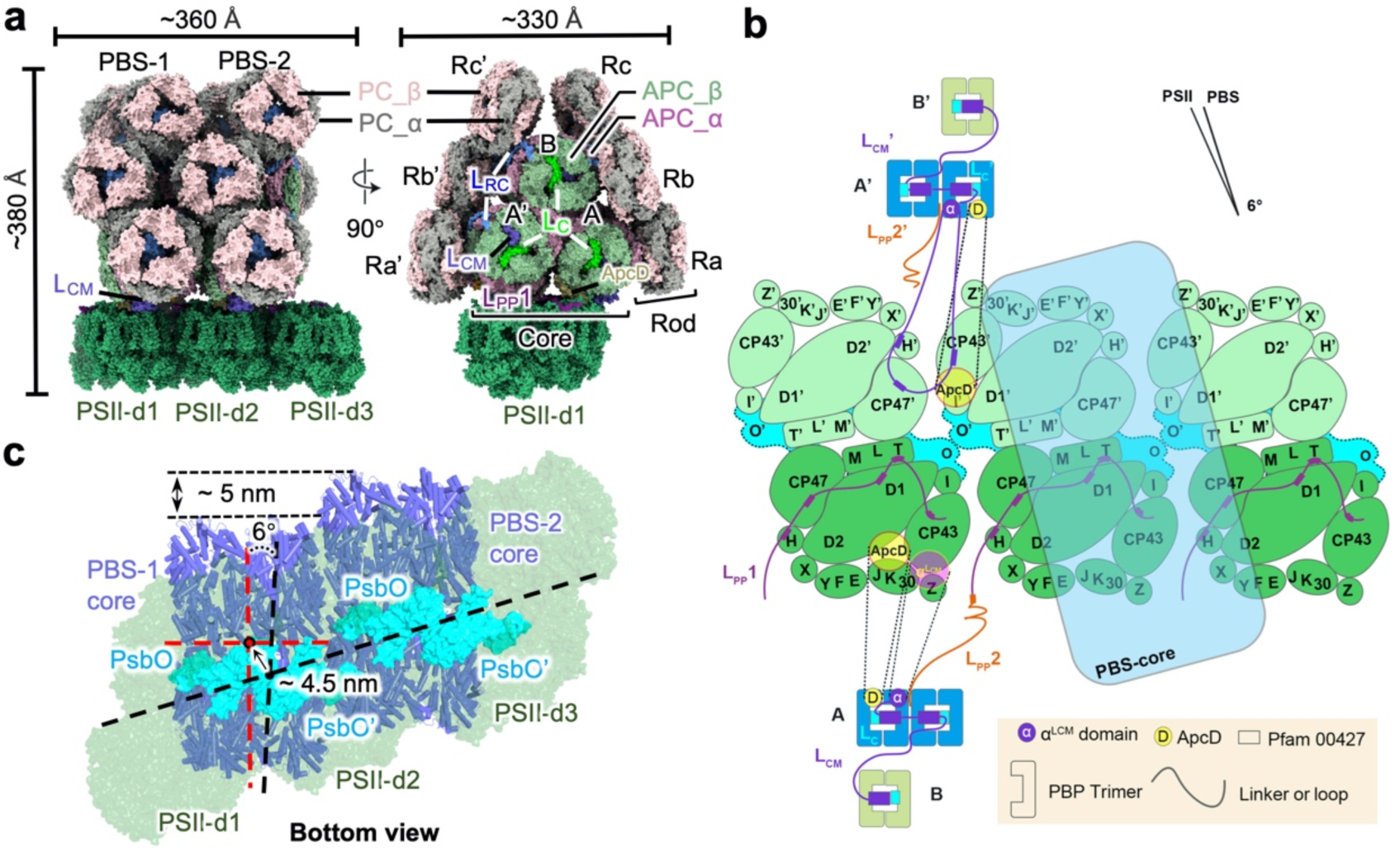
Overall structure of PBS-PSII supercomplex from *Spirulina platensis*. **a**, Overall structure of cyanobacterial PBS-PSII is shown as surface representation. The key subunits are color-coded as indicated. **b**, Schematic model of the PBS-PSII architecture, showing the components and connections of PBS and PSII. Transmembrane subunits are shown in PSII dimers. PSII dimer array displays the interaction between adjacent PSII dimers through PsbO. PSII dimers 1&2 display the interactions between PBS and PSII mediated by L_PP_1, L_PP_2, L_CM_-PB loop and ApcD. **c**, The bottom view of the PBS–PSII supercomplex shows that the PBS core plane rotates approximately 6° along the face plane of the PSII dimers. The centers of the PBS core and PSII-d1 and -d2 shift approximately 4.5 nm. The PBS-1 and -2 shift approximately 5 nm.

To analyze the spatial distribution of the PBS-PSII supercomplex on the native thylakoid membrane, the subtomogram maps were placed back into the original tomograms. Unlike red algae where PBSs are closely sandwiched between parallel thylakoid membranes and adjacent PBSs anchor to the upper and lower membranes in opposite orientations^20,24^, there is sufficient free space between the cyanobacterial PBS-PSII arrays, providing convenience for the movement of the PBS-PSII arrays along the membrane (Extended Data Fig. 3a-e). In addition, the 77 K spectra shows a significant increase in PSI fluorescence at 730 nm under orange light and dark adaptation compared with white light, suggesting that there might be a considerable number of PBSs close to PSI^25^ (described in Methods) (Extended Data Fig. 3f). Unfortunately, we could only obtain the *in situ* PBS-PSII structure without PSI associated. This situation is significantly different from that of red algae, where two PSI subcomplexes are connected to two ends of PSII array^20^, suggesting the interaction fashion of PSII with PSI is distinguished between cyanobacteria and red algae. This is likely reasonable because AFM observation for thylakoid membranes of cyanobacterium *Synechococcus elongatus* UTEX 2973 corroborated that PSII arrays are interspersed with PSI, which exhibits diverse interaction patterns and inhomogeneous arrangement in native thylakoid membranes^26,27^. We also identified 180 phycocyanobilin (PCB) for each PBS, and 70 chlorophylls, 4 heme molecules, 18 carotenoids and 56 lipids for each PSII dimer. All pigments are shown in Supplementary Table 1.

### *In situ* structure of cyanobacterial PBS and super-PBS

The *in situ* PBS structure of *S. platensis* has a typical hemi-discoidal morphology with a triangular core consisting of a top cylinder B and two basal cylinders A and A’ and six phycocyanin (PC) rods surrounding the core (Fig. 2a), which is consistent with the published cryo-EM structures of PBS from other cyanobacterial species^7,23^. However, the densities of rods, especially the peripheral rod hexamers, cannot be solved well because cyanobacterial rods are mobile and can switch conformation^28^, implying that cyanobacterial PBS may require broad space to accommodate the structural mobility (Extended Data Fig. 4a and b). There is only one type of rod-core linker protein (L_RC_) for the PBS of *S. platensis*. L_RC_ contains a conserved N-terminal Pfam00427 domain located in the cavity of the PC hexamer proximal to the core, and a C-terminal domain (CTD) protruding out of the central cavity of rod hexamer and latching onto the lateral grooves formed by α and β subunits of core cylinders forming several binding belts, similar to that in the published single-particle cyanobacterial PBS structures^7,23^ (Extended Data Fig. 5a-c). In addition, the *in situ* structure reveals that a short tail of L_RC_ CTD (R230 to L240) extends from the lateral grooves to the front surface, serving as a buckle to stabilize and strengthen the stacking of PBS core layers, which may contribute to energy migration between adjacent PBSs (described below) (Extended Data Fig. 5d).

In cyanobacteria *S. platensis*, PBSs are arranged into linear rows on the stromal side of PSII with the cores stacked tightly in a face-to-face manner (Fig. 2a). However, due to the approximately 5 nm shift between two adjacent PBS cores (Fig. 2c) and the slight twist (∼ 25.3°) between two basal cylinders (Extended Data Fig. 4c and d), the hexamers from adjacent cores are positioned with corresponding displacements.

Interestingly, the offset of 5 nm is about the same as the radius of the hexameric ring, which ensures that there are still certain interfaces between the two adjacent cores. In detail, the interactions between the core layers B1’/A1/A4’ and B1/A4/A1’ are formed by several hydrogen bonds (H-bonds) between the outermost trimeric β layers (Extended Data Fig. 6). Moreover, cyanobacterial L_CM_ has evolved an extra loop region (D527 to K550) extending to the central cavity of core A1 layer (Extended Data Fig. 7a-c). We found that this loop bound to A1 layer and L_C_ via 7 H-bonds (D527 – R107/β2_A1_, D527 – Q23/L_C_, S535 – T28/L_C_, N540 – D105/β2_A1_, K541 – S118/β1_A1_, N544 – Q2/β2_A1_ and G546 – R77/β1_A1_), which further stabilizes the conformation of A1 layer (Extended Data Fig. 7a-b). The binding belts and the short buckles formed by L_RC_ CTDs are also involved in the compact assembly of PBS (as mentioned above) (Extended Data Fig. 5b). All these lead to the dense face-to-face packing of PBSs with the bilin distances between adjacent PBSs in~30 Å (Extended Data Fig. 6b). Such a tightly stacked arrangement of PBSs may function as super-PBS in which the excitation energy flow could propagate across the adjacent PBSs along the PBS rows (described below).

### The binding mechanism of PBS with PSII in cyanobacteria

Efficient excitation energy transfer from the PBS to PSII depends on the accurate interactions between PBS terminal emitters (ApcD and L_CM_) and PSII^11^. In the *in situ* PBS-PSII supercomplex structure, we clearly resolved the loop of the phycobilin-binding domains (PB domain) of L_CM_ (PB-loop/L_CM_), which was absent in the published *in vitro* PBS structures (Extended Data Fig. 7d and e).

As mentioned above, L_CM_ and L_CM_’ have different interaction manners with PSII. L_CM_ interacts with PSII-d1 via α^LCM^ rather than through the PB-loop (Fig. 3a-c), while L_CM_’ uses PB-loop’ to contact with both PSII-d1 and PSII-d2 through H-bond interactions of D118, N119 and D134 of PB-loop’ with R476/PSII-d1_CP47’_, K227/PSII-d1_CP47’_ and Y130/PSII-d2_CP43’_, and through T-shaped π-π interaction of its aromatic residue F132 with F133/PSII-d2_CP43’_ (Fig. 3d and e). The sequence alignment of PB-loop in different cyanobacteria shows that the key residues D118 and F132 maintain the corresponding amino acid properties, indicating that PB-loop may mediate the association between PBS and PSII via some key residues in cyanobacteria (Extended Data Fig. 8a). Similarly, ApcD and ApcD’ have different contacts with PSII dimer. ApcD interacts with D2 of PSII-d1 through several electrostatic interactions, while ApcD’ binds with PSII dimer through a H-bond (Fig. 3f and g). Sequence alignment of ApcD shows that these charged residues are basically conserved in different cyanobacteria, suggesting that ApcD from different cyanobacteria may contact with PSII in a similar interaction manner (Extended Data Fig. 8b).

**Fig. 3.**
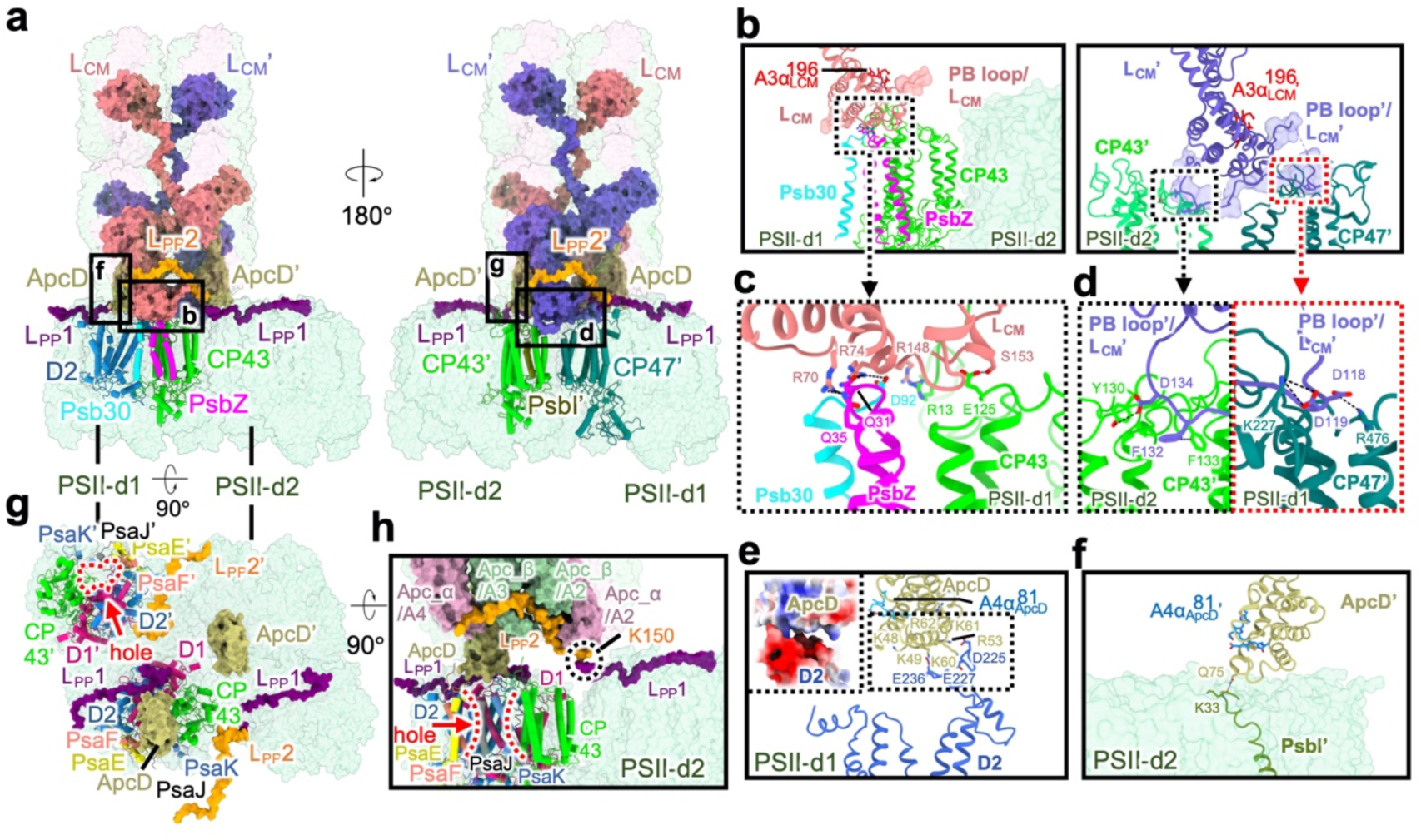
Interactions of PBS with PSII tetramers. **a,** Different views of interactions between PBS and PSII tetramers. L_CM_ (L_CM_’), ApcD (ApcD’) and L_PP_2 (L_PP_2’) of PBS and L_PP_1 of PSII dimers are highlighted. The transmembrane subunits involved in the interactions are shown as cartoon. **b,** Enlarged views showing L_CM_ binds to only PSII-d1 (left panel), while L_CM_’ binds to both PSII-d1 and -d2 (right panel). PB-loop/L_CM_ (PB-loop’/L_CM_’) is shown as surface representation in 60% transparency. **c, d** Details of the interactions of L_CM_ (**c**) and L_CM_’ (**d**) binding to PSII subunits. The residues involved in the interactions are shown as stick representation. **e, f**, Interactions of ApcD (**e**) and ApcD’ (**f**) with PSII dimers. The enlarged view of (**e**) showing the electrostatic interaction between ApcD and D2 subunit of PSII-d1. **g, h**, different views showing the position of L_PP_2 and the interaction between L_PP_2 and L_PP_1 of PSII-d2. The positively charged residue K150 of L_PP_2 is labeled.

In addition to L_CM_ and ApcD, a homologue of linker 2 of PBS and PSII (L_PP_2) of the red algal PBS (Ref) was found in the *in situ* structure of the *S. platensis* PBS, and also named as L_PP_2. Using its C-terminus L_PP_2 has a slight contact with the N-terminus of the linker 1 of PBS and PSII (L_PP_1), a new component of PSII dimers (discussed later), via a conserved positively charged residue K150 (Fig. 3h and i, and Extended Data Fig. 9e-g). Given that the complete L_PP_2 has not been solved yet, it could not be excluded that its N-terminal region has a direct interaction with PSII. However, even if this interaction exists, it may be transient and unstable in *S. platensis*, since we cannot obtain the structure. By contrast, the red algal L_PP_2 forms a stable interaction with the CNT subunit of PSII dimer directly by its N-terminal loop, exhibiting a much stronger binding between PBS and PSII than cyanobacteria^20^. Since the interaction forces between PBS and PSII are mainly attributed to several polar interactions provided by loop regions of the PBS-PSII interface, and no complete linker ties PBS and PSII together, the coupling between cyanobacterial PBS and PSII seems weak and unstable *in vivo*^29,30^. It is quite different from red algal PBS-PSII, where PBS is tightly bonded on the PSII surface by four PBS-PSII linker proteins (L_RC_2, L_RC_3, L_PP_2, L_PP_1), as well as L_CM_ and ApcD^20^. This may provide a structural basis for the diffusion movement of cyanobacterial PBS on the surface of thylakoid membrane observed by fluorescence spectra and fluorescence recovery after photobleaching (FRAP) measurements^29,31^.

### *In situ* structure of cyanobacterial PSII and interaction pattern

Similar to published cryo-EM structure of cyanobacterial PSII^32^, the *in situ* structure of *S. platensis* PSII monomer consists of 17 core subunits (D1, CP47, CP43, D2, PsbE, PsbF, PsbH, PsbI, PsbJ, PsbK, PsbL, PsbM, PsbT, PsbX, PsbY, Psb30 and PsbZ) and 4 extrinsic subunits (PsbQ, PsbO, PsbU and PsbV) (Supplementary Table 1). Notably, we observed an unprecedented density that attached on the stromal side of PSII (Fig. 4a). Because the PSII-binding position of this density is similar to that of the red algal L_PP_1^20^, it may be a homolog of the red algal L_PP_1 and also named L_PP_1 here. Unfortunately, the sequence of L_PP_1 could not be identified in our structure, so it is modeled as poly-Ala residues. L_PP_1 contains an N-terminal L_PP_2-binding motif (A1 to A9) followed by a PSII-binding motif (A10 to A47), which attaches to the PSII surface through binding with D1, D2, CP43, CP47, PsbT and PsbH by electrostatic interactions (Fig. 4b, Extended Data Fig. 10a - c). Remarkably, only one monomer of the PSII dimer has the L_PP_1 protein attached, as the corresponding site for L_PP_1 binding on the other monomer is occupied by a segment of the PB-loop/ L_CM_. Actually, this segment shares a highly similar structural feature and the PSII binding pattern with L_PP_1 (Extended Data Fig. 10d). Notably, the C-terminus of the red algal L_PP_1 (PBS-core-binding motif) is embedded into the groove of L_CM_ to tie PBS and PSII tightly^20^, while only PSII-binding motif is resolved in cyanobacterial L_PP_1. This is because the outmost APC_β trimers of cylinder core A /A’ occupy the binding site, therefore the C-terminus of cyanobacterial L_PP_1 may be instable and unresolved (Fig. 2a).

**Fig. 4.**
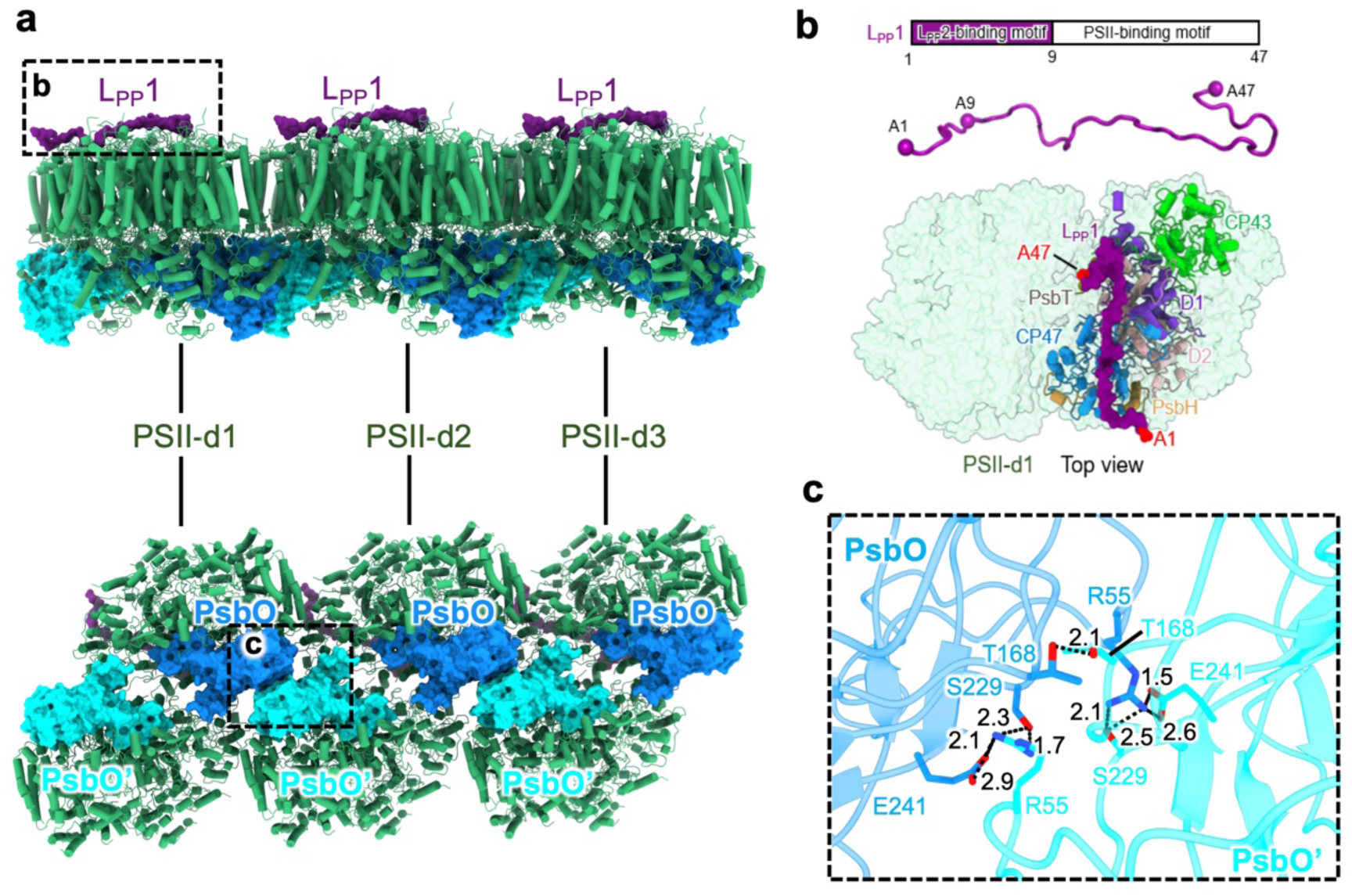
Interaction pattern between PSII dimers. **a**, The interaction pattern of PSII dimers from the side view (upper) and the bottom view (lower). L_PP_1, PsbO and PsbO’ are shown as surface representation and colored in purple, dodger blue and cyan, respectively. **b**, The diagram of structural element of L_PP_1(upper panel) is shown above the atomic model (middle panel). Lower panel: Top view of the stromal side of PSII showing that L_PP_1 spans on the stromal surface of one PSII monomer created by transmembrane helices of D1, D2, CP43, CP47, PsbH and PsbT. The N- and C-terminal residues of L_PP_1 are colored in red. **c**, Close-up view of the interaction between PsbO molecules from adjacent PSII dimers. The residues involved in the interaction are shown as stick representation.

In addition, structural analysis shows that the connection between PSII dimers only via two PsbO subunits on the lumen side (Fig. 4a). Two conserved charged residues R55 and E241 of PsbO interact with E241’ and R55’ of PsbO’ forming salt bridges (Fig. 4c and Extended Data Fig. 8c). Besides, two conserved polar residues T168/ PSII-d1_PsbO’_ and S229/ PSII-d1_PsbO’_ are also contributed to the connection of PSII dimers through forming H-bonds with T168/ PSII-d2_PsbO_ and R55/ PSII-d2_PsbO_ (Fig. 4c). Cyanobacterial PSIIs are virtually stably arranged in long parallel rows *in vivo*, therefore PsbO mediated interactions seem to be the crucial factors for the array layout of PSII dimers.

### Multiple “super-PBS” mediated collaborative energy transfer pattern

As mentioned above, cyanobacterial PBSs are stacked closely with each other, forming a unique “super-PBS” organization, in which the excitation energy flow could propagate across the adjacent PBSs (Fig. 2a). Several bilin pairs of the super-PBS, having the short distances between two neighboring PBS layers, are probably key bridges in the energy transfer across the interface (Extended Data Fig. 6b). It should be noted that terminal emitters (^A4^α_ApcD_/^A4^α_ApcD_’ and ^A3^α_LCM_/^A3^α_LCM_’) are located in the layer A3/A3’ and A4/A4’ of PBS core, thus the energy from layer A1/A1’ will have to travel across layer A2/A2’ to reach the terminal emitters within one PBS core. However, in the super-PBS the layer A1/A1’ from one PBS becomes neighbor to the layer A4/A4’ from the other PBS, which could deliver the energy from layer A1/A1’ directly to the terminal emitters ^A4^α_ApcD_/^A4^α_ApcD_’ of neighbor PBS. Based on these data, we speculate that the cyanobacterial “super-PBS” might increase the energy transfer efficiency by coupling pigments of adjacent PBSs.

Terminal emitters of PBS could converge the energy absorbed by PBS and further transfer it to PSII^8,11,33^. Owing to the rotation and shift between PBS and PSII dimers (as mentioned above), the distances from four terminal emitters, ^A4^α_ApcD_, ^A4^α_ApcD_’, ^A3^α_LCM_ and ^A3^α_LCM_’, to the nearest chlorophylls in PSII are different, which leads to an asymmetry in the energy distribution to two PSII monomers (Fig. 5a). Similar to red algal PBS-PSII supercomplex, several aggregated chlorophylls of CP43 (CP43’) and CP47 (CP47’) form the displaced-parallel Chls clusters, prompting the delocalization of π system and downgrade the energy level^20^ (Fig. 5b and c). They are considered as the key mediators of energy transfer from PBS to P680^34^ (Fig. 5a). Notably, the energy could be transferred from adjacent PBSs to PSII dimers due to the formation of the “super-PBS” array. Taking PSII-d2 as an example, ^A4^α_ApcD_, ^A3^α_LCM_ and ^A3^α_LCM_’ of PBS-1 could also deliver the energy to P680’ and P680 of PSII-d2, therefore each P680 could accept the excitation energy from two adjacent PBSs (PBS-1 and PBS-2) (Fig. 5a).

**Fig. 5.**
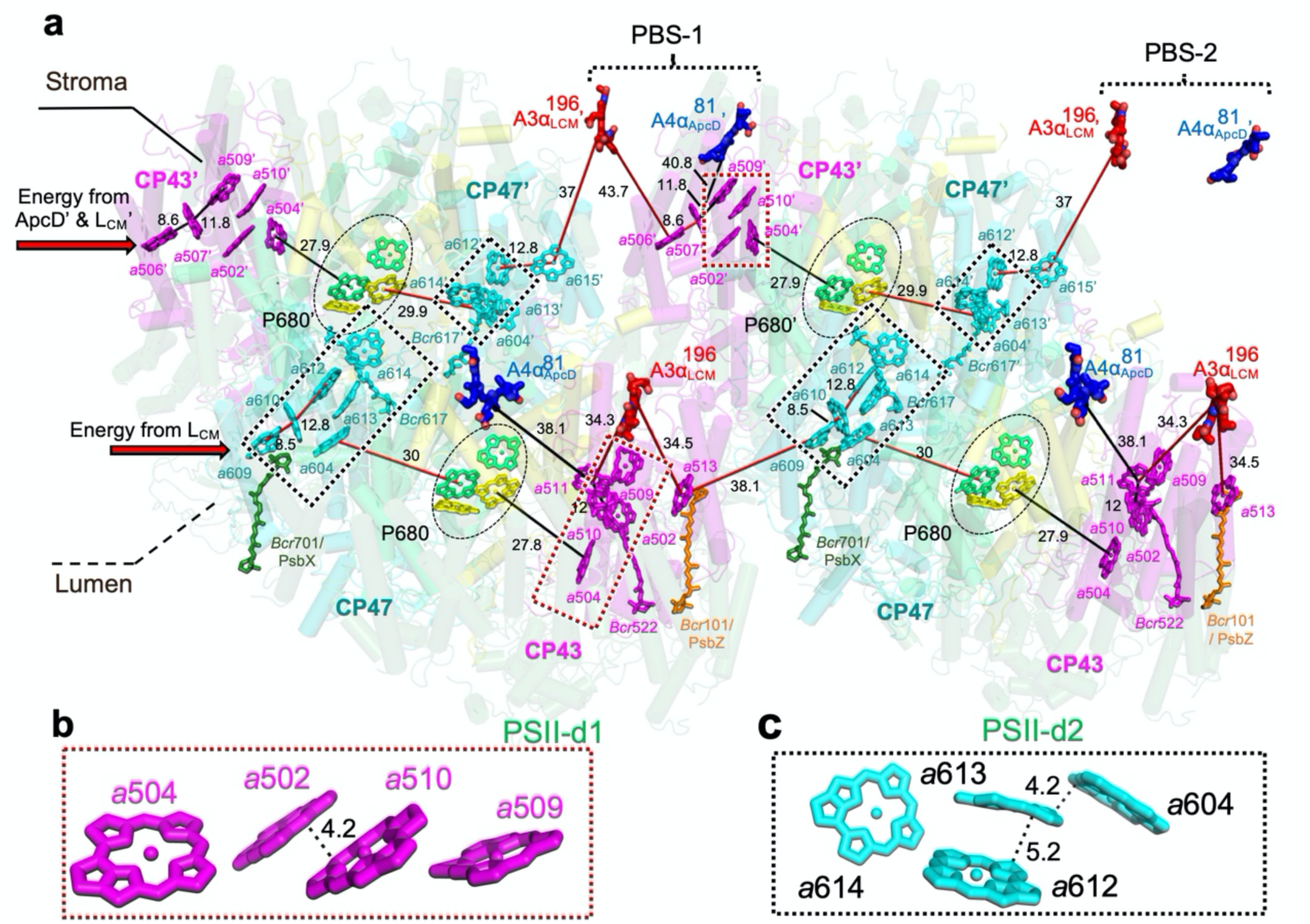
Key pigment arrangements and possible energy transfer pathways. **a**, Distribution of key pigments and possible energy transfer pathways in PBS-PSII supercomplex from the top view normal to the membrane plane. The key pigments are shown as bold-stick and the π-π distances (Å) for the adjacent pigments are labeled in black. P680 are boxed in oval. The low energy state Chl clusters of PSII are boxed in rectangle. The red arrows indicate the direction of the potential energy transfer from ^A3^α_LCM_, ^A3^α_LCM_’ and ^A4^α_ApcD_’ from neighboring PBSs. **b**, **c**, Magnified views of the low energy state Chl clusters of CP43 (**b**) and CP47 (**c**).

In conclusion, our STAgSPA, combining the advantages of cryo-ET-STA and SPA, proposes a novel *in situ* structural analysis strategy. This strategy has been successfully applied to the *in situ* structural determination of photosystems in both red algae^20^ and cyanobacteria, providing a new methodological paradigm for improving the *in situ* structural resolution of cellular and tissue samples. In addition, the *in situ* high-resolution structures of the PBS-PSII supercomplex from cyanobacteria and red algae revealed remarkable differences in the association pattern and energy transfer mechanism of PBS-photosystems between these two photosynthetic organisms: (1) in red alga *P. purpureum*, four PBS-PSII linker proteins mediated the tight association between PBS and PSII, while the cyanobacterial PBS-PSII exhibits loose and instable binding pattern due to the lack of the similar linkers; (2) in red algae, two PSIs are stably associated at both ends of PSII arrays that stacked by two or four PSII dimers, while cyanobacteria form the longer PSII arrays that may lack the stable association with PSI. Thus, we can only obtain the stable *in situ* structure of PBS with PSII without PSI resolved; (3) there are no interactions between adjacent PBSs in red algae, while cyanobacterial PBSs are densely stacked together, forming the “super-PBS” pattern. Thus, the PSII reaction centers of cyanobacteria could accept the excitation energy from multiple adjacent PBSs, forming the sophisticated pigment networks for the efficient energy transfer from PBS to PSII, which could effectively compensate for the disadvantage of fewer bilins in cyanobacterial PBS compared to red algae^20^.

## Methods

### Strains and culture conditions

*Spirulina platensis* FACHB-439 was bought from Freshwater Algae Culture Collection at the Institute of Hydrobiology, FACHB. Cells were grown in *Spirulina* medium at 25 °C and shaken at 150 rpm under continuous low orange light illumination with an intensity of ∼ 10 μmol of photons m^-2^ s^-1^, which was provided by LED with wavelength at 625 nm^35^. The cells harvested at the middle-logarithmic phase of growth were used for experiments.

### 77k fluorescence emission spectra

*Spirulina platensis* FACHB-439 cultured as described above was collected and then kept in the dark for 20 minutes before measuring fluorescence spectroscopy. Fluorescence emission spectra was recorded by FS5 (EDINBURGH) at 77 K with an excitation wavelength of 580 nm. Cells cultured under low white light (∼ 10 μmol of photons m^-2^ s^-1^) were used as the control trial.

### Cryo-EM sample preparation and cryo-FIB milling

Cells were preprocessed in the same way as for 77k fluorescence emission experiments before freezing. 4 μl suspended cells were loaded onto the front side of glow-discharged holy-carbon copper grids (Quantifoil R1.2/1.3, 200 mesh), and 2 μl medium was subsequently added to the back side of the grids. The grids were back-side blotted by Leica EM GP2 (Leica Company) for 6 seconds at 20 °C and 100 % humidity, and plunged into liquid ethane at -184°C for vitrification. Grids were stored in liquid nitrogen until used for FIB milling. Cryo-FIB milling was performed using an FIB/SEM dual-beam microscope (Aquilos, Thermo Fisher Scientific). Briefly, the cryo-EM grids were clipped into cryo-FIB Autogrids in the liquid nitrogen reservoir, and loaded into the FIB/SEM chamber at the high vacuum and -190 °C. To reduce sample charging and protect the sample, the surface of samples was sputter coated with a layer of platinum and then deposited an organometallic platinum using gas-injection system (GIS) operated with a 35 s gas injection time before milling at a work distance of 11 mm. The stage was tilted at 22° (milling angle of ∼ 15° related to the EM grid). Micro-expansion joints were milled to release stress in the support film^36^. Rough milling was performed with the accelerating voltage of the ion beam at 30 kV and a current of 3.0 nA to 0.5 nA to produce lamellae with a thickness of ∼ 500 nm. Due to the inherent Gaussian profile of the ion beam, the typical milling scheme causes preferential thinning of the top of the lamella, resulting in uneven thickness of the lamella, therefore an optimization of the milling technique was used in the further polishing step^37^. In brief, rough-milled lamellae were polished by tilting each side of the lamella 1° towards the ion beam and milling with 50 – 30 pA. The optimized milling procedure results in final lamellas with homogenous thickness (100 – 200 nm) across their entire length.

### Cryo-ET data collection and tomogram reconstruction

The data collection of the prepared lamellas was performed on a 300 kV Titan Krios EM G3i (Thermo Fisher). TEM images of lamellas at low and media magnifications were presented in Extended Data Fig.1a and b. Before data collection, the stage was tilted about +15°/-15° to make the lamella plane approximately perpendicular to electron beam. Tilt series were acquired by a K3 Summit direct detector camera (Gatan) at magnification of 53,000 × (equivalent pixel size 1.632 Å) with a post-column Quantum energy filter (Gatan) in zero-loss mode and slid width of 20 eV. Each tilt image was recorded as a super-resolution movie stack containing 15 frames with a total dose of 4.4 e^-^/Å^2^. Software SerialEM^38^ and dose symmetry tilting scheme were used to collect the data. The tilted images were collected with angular range from - 42° to + 42° (related to lamella plane) and an interval step of 3°. Each tilt series contains 29 images with a total dose of ∼130 e^-^/Å^2^. The defocus range used in data collection was from -2.5 μm to -5.5 μm.

The movie stacks were corrected for the beam-induced motion by using MotionCor2^39^ and summed into images with a binned factor of 2. Images of tilt series were then merged into stacks and aligned by patch tracking in IMOD^40^. In total, ∼200 tomograms were reconstructed, and 94 tomograms with good alignment were selected out for subsequent analysis. Tomograms were 3D reconstructed at bin 8 in IMOD^40^ by Weighted Back-Projection method and processed with a deconvolution script ‘tomo_deconv’ (https://github.com/dtegunov/tom_deconv) before segmentation and particles annotation.

### Subtomogram averaging of PBS complexes

The reconstructed tomogram showed *S. platensis* sample with a regular distribution of PBS particles (Extended Data Fig. 1c). Initially, ∼3000 PBS sub-tomograms were manually picked from 6 tomograms (at bin 8) in Dynamo^41^. After translational and rotational alignment in Dynamo at bin 4, the averaging results showed a shape of PBS array at ∼ 30 Å resolution. By use this density map as template, ∼60,000 particles were picked out by template matching from 94 selected tomograms in emClarity^42^. After manually removed the particles of falsely pickings, 53,000 particles were subjected for subtomogram averaging. The cycles of alignment and averaging were consecutively processed at bin 4, bin 3 and bin 2 with a shape mask containing three PBSs (mask does not include the membrane region and the peripheral hexamer of each rod), yielding an averaging map of ∼ 10 Å. The masked density map was showed in Extended Data Fig. 1d. The unmasked density map had a blurred density at the bottom region of PBS, which were assumed to be PSII signal in thylakoid membrane.

### Tomogram segmentation and 3D visualization

The thylakoid membrane in tomograms were segmented by using TomoSegMemTV^43^ and manually tracing the membrane density in the regions with weak signal. PBSs were repositioned back to tomograms according to the translational coordinates and Euler angles from refinement result of subtomogram averaging. PSII particles were simply posited according to the spatial relation to PBS particles. PBS array, PSII and the thylakoid membrane were represented as different colors in ChimeraX^44^ (Extended Data Fig.3b, d and f).

### Data collection for high-dose single-particle images

Micrographs were acquired as 80 movie frames with a total dose of 35 e^-^/Å^2^ by using SerialEM^38^ at same imaging condition as the tilt series data with a defocus range from -1.5 μm to -4.0 μm. Beam induced motions between frames were corrected in cryoSPARC^45^ and the output micrographs were dose-weighted and binned with a factor of 2. The contrast transfer function (CTF) parameters of micrographs were estimated by CTFFIND4^46^. Micrographs were inspected visually to exclude very large motion or incorrect CTF estimation. 1652 micrographs were selected from ∼2200 micrographs for further analysis.

### Particle picking from single-particle images

To translationally and rotationally localize the PBS-PSII complexes in high-dose images of lamella sample, we performed high-resolution template matching method using a recently developed software *is*SPA^15^, which was optimized to identify target protein signals from overlapping densities of crowded environment. The result density map from subtomogram averaging was applied with a bandpass filter from 10 Å to 50 Å and a mask containing three PBSs and was used as the 3D template. Sampling points of three Euler angles were defined by dividing the Euler angle space equally at an interval of 3.7° in RELION^47^ Euler angle definition. The cross-correlograms were calculated at bin 2 level and the threshold used for particle detection was set as 6.3. Reduplicated peaks in cross-correlograms within a distance of 15 pixels and an orientation spacing angle of 30° were exclude. An average of ∼130 peaks were detected in each micrograph (an example of detected particles was demonstrated in Extended Data Fig. 1e). Subsequently, the parameters of each particle, including coordinates, defocus and Euler angles were imported into RELION^47^ for structure-refinement. The first attempt of particle picking generates 212,000 detections. By using the averaging map of sub-tomograms as the initial reference, refinement in RELION^47^ yielded a density map of 6.5 Å (Extended Data Fig.1f). This density map was used to preform high-resolution template matching once more. The second time particle picking detected 224,000 particles. All the particles were merged together and the overlapped particle were removed. The remaining 167,000 particles were subjected to further analysis.

Although the membrane region was excluded in high-resolution template matching by applying a mask, the reconstruction from 2D particles picked by *is*SPA^15^ showed a clear PSII signal. This signal could also be identified from raw tomograms (Extended Data Fig.3a, c and e), which indicated that the structure was not induced by template bias.

### Structure-refinement on PBS-PSII complexes

The detailed steps of structure-refinement for subregions of PBS-PSII complexes were represented in Extended Data Fig. 2. Particles were shifted to the center of two PBS monomer and extracted at bin1 with a box size of 400 pixels. A local refinement (with angle step of 3.7° and a mask containing two PBS along with a membrane region) yielded a 5.6 Å reconstruction. Particles was subjected to three rounds of CTF refinement and 3D auto-refine to increase the defocus estimation accuracy. Then, 3D classification was carried out with a smaller mask focusing on the PSII region to filter out falsely picked particles and low-quality particles. This 3D classification yielded 4 output classes, while the first two classes showed clear PSII density and a same conformation. 127,000 particles from class 1 and class 2 (rotated 180° around *z* axis comparing to class 1) were selected to reconstruct a lower resolution (5.1Å) PBS-PSII array density map (refined with a mask containing 2 PBS) and subjected to further process.

In cryo-ET results, PBSs were distributed in arrays bent into different curvatures, and densely packed and overlapped in *z* direction. The high-resolution template matching step using the reference of three PBS density might miss potential particles. For samples of 1D protein assembly, more particle positions could be inferred from those already detected by extending the coordinates^48^. In order to pick the particles exhaustively, a refinement with a mask containing two PBS APC was performed and particles were subsequently shift-expanded to 1 left position (shifting the x, y in reconstruction coordinates by -15, -70 pixels, while keeping the Euler angle unchanged) and 1 right position (shifting by 15, 70 pixels). One step of local refinement was performed to align translations and rotations of these particles, and overlapped particles within 30 Å were removed (270,000 particle remained). Then the particles were further expanded to 3 left positions (-15, -70 pixels, -30, -140 pixels, -45, -210 pixels) and 3 right positions (15, 70 pixels, 30, 140 pixels, 45, 210 pixels), increasing the total particles number to 1,873,000. Again, the particles were aligned and the overlap particles were removed (1,179,000 particle remained). A mask focusing on a smaller subregion containing only one APC was used for the following 3D classification, and 664,000 particles were selected to further processing.

To improve the resolution of the APC cores, the particles were subjected to an additional 3D classification in RELION^47^ and a local refinement in cryoSPARC^45^. Then we obtained a 3.6 Å local structure. The similar processing steps were also applied on the PSII region. Two steps of refinements were performed on selected 83,000 particles, first with a shape masked containing 2 PSII dimer and then with separate local masks on each PSII dimer. Local refinements in cryoSPARC^45^ improved the resolution of each dimer of PSII to 3.5 Å and 3.6 Å respectively. All the resolutions were estimated by gold standard Fourier Shell Correlation with a criterion of 0.143 (Extended Data Fig. 2e-j).

Since there was no specific difference found between the two connected PSII dimer, we suggest a periodic symmetry in the structure of PBS-PSII complexes. To better display the whole complexes, 2 copies of regional map of APC (3.6Å) and 3 copies of regional map of PSII (3.5Å) were fitted into a lower resolution (5.1Å) PBS-PSII array density map and merged as one.

### Model building and refinement

For model building of the PBS, the atomic model of *Synechococcus* 7002 PBS (PDB entry 7EXT)^33^ was first docked into the maps using Chimera^49^. The peripheral PC hexamers were not well-fitted into the maps, and thus were manually deleted. Then the PBP and linker proteins were mutated to the corresponding sequences of *S. platensis* in Coot^50^ and manually adjusted to better fit with the map. The sequence of L_PP_2 was identified by searching the homologue of sll1873 (ApcG) in *Synechocystis* 6803^7^ based on the Basic Local Alignment Search Tool (BLAST). Sequence assignments of PB-loop of L_CM_, L_RC_ and L_PP_2 were guided by corresponding residues in Coot^50^. Furthermore, *de novo* model building was performed on the unidentified subunits defined as the L_PP_1, and the N/C terminus were artificially defined in this study. L_PP_1 is located on the stromal side of PSII dimers. However, no similar structure was observed in published PSII structures, indicating that it might be lost during purification. To build the model of PSII dimers, the structure of *Synechocystis* 6803 (PDB entry 7RCV)^9^ was docked into density maps using Chimera^49^ and the sequences were mutated to those of *S. platensis* with Coot^50^.

The model buildings of PBS and PSII dimers were completed via iterative rounds of manual building with Coot^50^ and refinement with phenix.real_space_refine^51^ with geometry and secondary structure restraints. Then, all parts were merged to the whole PBS-PSII supercomplex and the structural validation report were produced against a whole artificially stitched map using local maps in MATLAB. The statistics for data collection and structure refinement are summarized in Extended Data Table 1. Figures were prepared with UCSF ChimeraX^44^ and PyMOL (http://pymol.org). The sequence alignments were performed by ClustalX2^52^ and created by ESPript^53^.

## Data Availability

The cryo-EM density map and atomic models have been deposited in the Electron Microscopy Data Bank and the Protein Data Bank for the PBS-PSII supercomplex structure (PDB ID code 8H4W), the whole artificially stitched map (EMDB ID code 34487), the local map of PBS at 3.5 Å resolution (EMDB ID code 34485), the local map of PSII dimer at 3.6 Å resolution (EMDB ID code 34486) and the low-resolution map of PBS-PSII at 5.1 Å resolution (EMDB ID code 34521).

## Acknowledgements

We thank the staff at the Tsinghua University Branch of the National Protein Science Facility (Beijing) for their technical support on the cryo-EM and the high-performance computation platforms. We thank Dr. Wei Lu from Department of Chemistry, Southern University of Science and Technology, for his assistance on 77K fluorescence spectra. This work was supported by the National Natural Science Foundation of China (32241030 and 32271245 to S.-F.S.), China Postdoctoral Science Foundation (2023M741985 to X.Y.), and the National Basic Research Program (2016YFA0501101 and 2017YFA0504600 to S.-F.S.).

## Author Contributions

S.-F.S. supervised the project; Y.X. and X.Y. frozen samples, performed cryo-FIB milling and the sequence analysis; X.Z. designed the cryo-EM strategy and optimized the data collection scripts; X.Z., X.Y. and Y.X. collected the cryo-EM data; X.Z. performed the cryo-EM analysis; X.Y. and Y.X. performed the model building and the structure refinement; Y.X. performed the biochemical experiments; X.Y., Y.X., X.Z., S.S. and S.-F.S. analyzed the structure; all authors contributed in writing the manuscript.

## Additional information

### Supplementary information

The online version contains supplementary material.

### Correspondence and requests for materials

should be addressed to Sen-Fang Sui.

