## Supplemental Information for "*In situ* structural determination of cyanobacterial phycobilisome-PSII supercomplex by STAgSPA strategy"

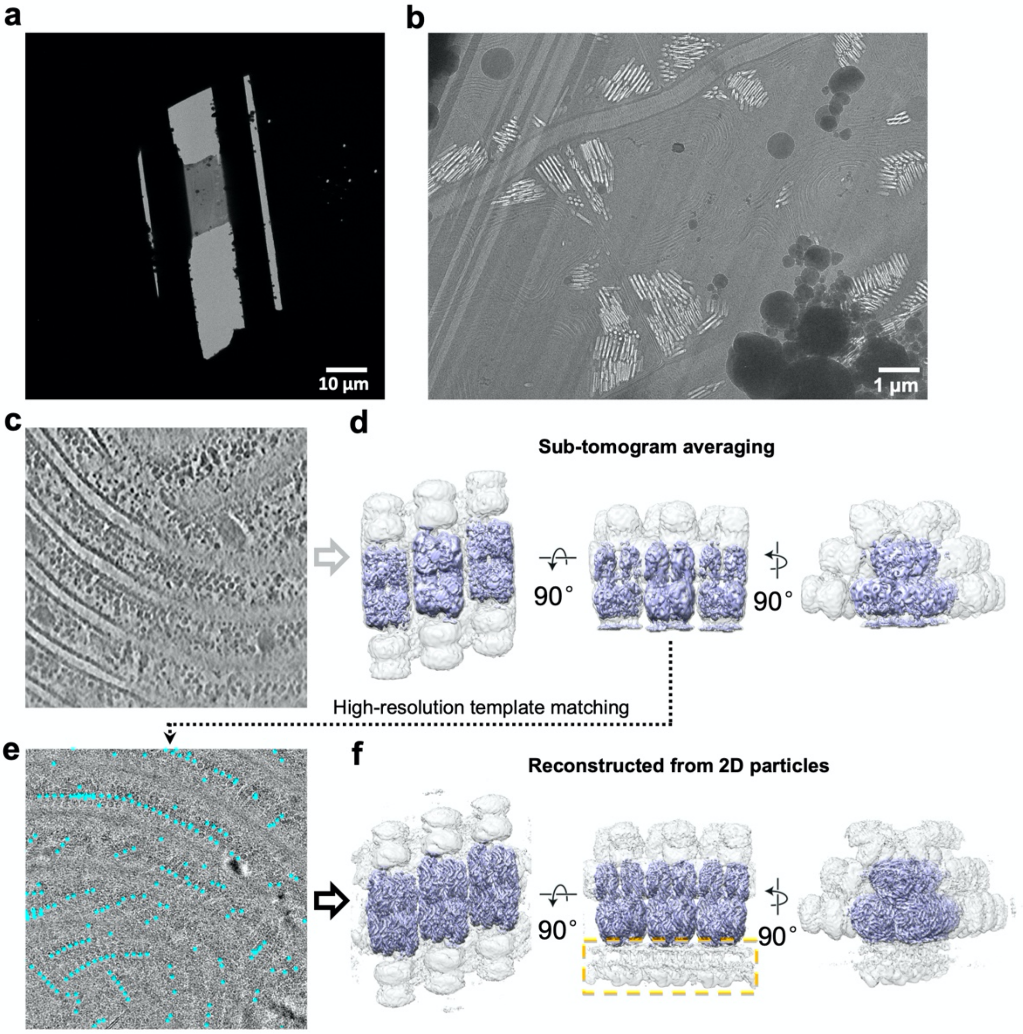


**Extended Data Figure 1 | Overview of FIB lamella sample preparation and flowchart of cryo-EM study.**

**a**, **b,** Representative EM images of *S. platensis* lamella (selected from 94) at low (**a**) and media (**b**) magnifications. **c**, A representative view of a Z-section of reconstructed tomogram (selected from 94 tomograms). **d**, Sub-tomogram averaging were processed with a mask only containing three PBSs, yielding a ~ 10 Å structure. The regions of APC cores are colored in purple. **e**, Particle picking from single-particle images by high-resolution template matching method using the result density map from sub-tomogram averaging as the 3D template. An example of detected 2D particles (cyan dots) of one micrograph (from ~2,200 images) was demonstrated. **f**, The reconstruction of 2D particles showing an increase in the overall resolution and extra density of the membrane region (yellow box).

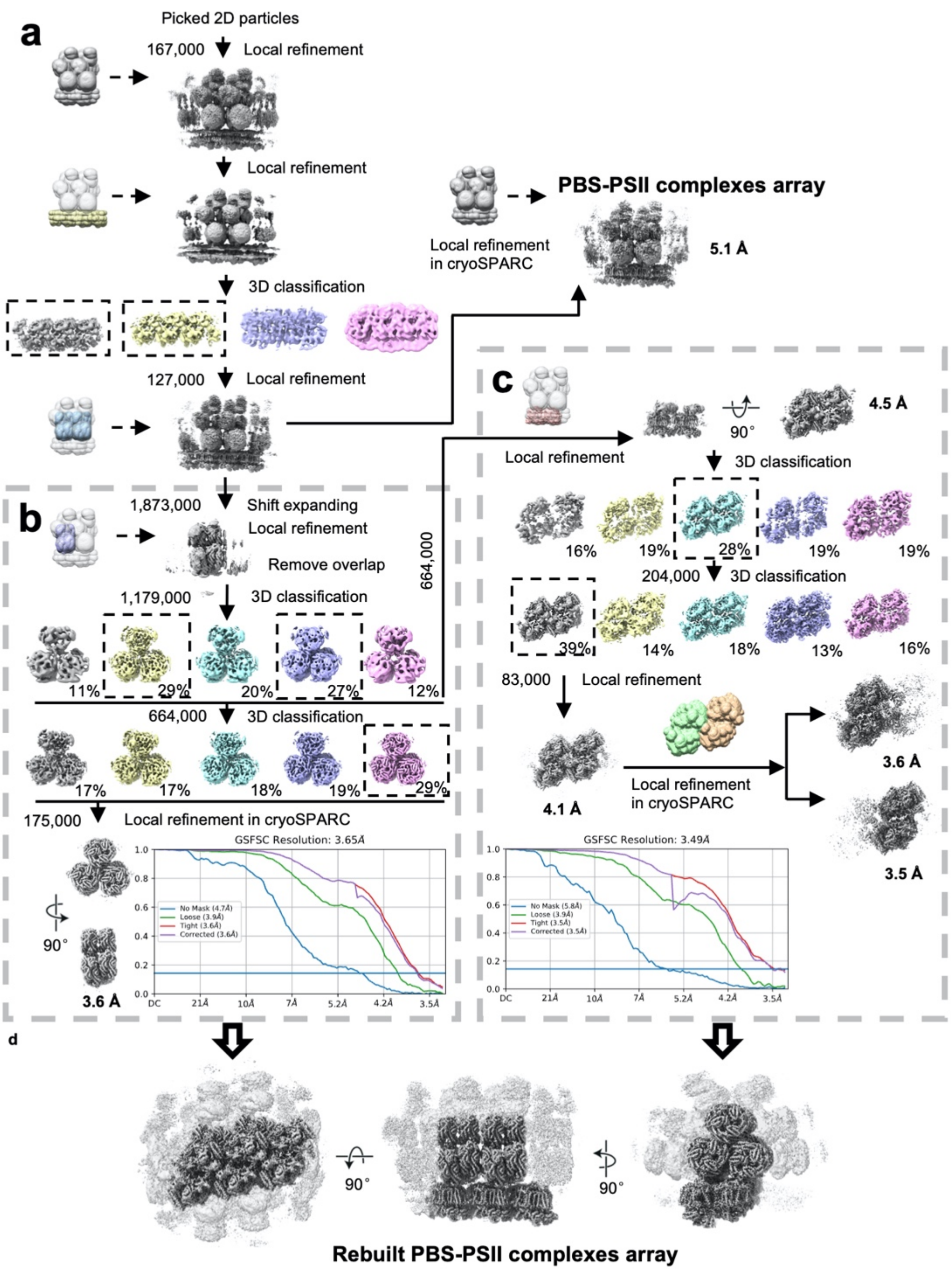


**Extended Data Figure 2 | Structure-refinement for subregions of PBS-PSII complex.**

**a**, Image processing procedure for the intact PBS-PSII complex. **b**, Image processing procedure for sub-region of PBS cores. **c**, Image processing procedure for PSII region. **d**, The rebuilt PBS-PSII complex array. Sub-regional structures of PBS cores and PSII dimers were merged as one artificial density map of PBS-PSII complex array in different views.

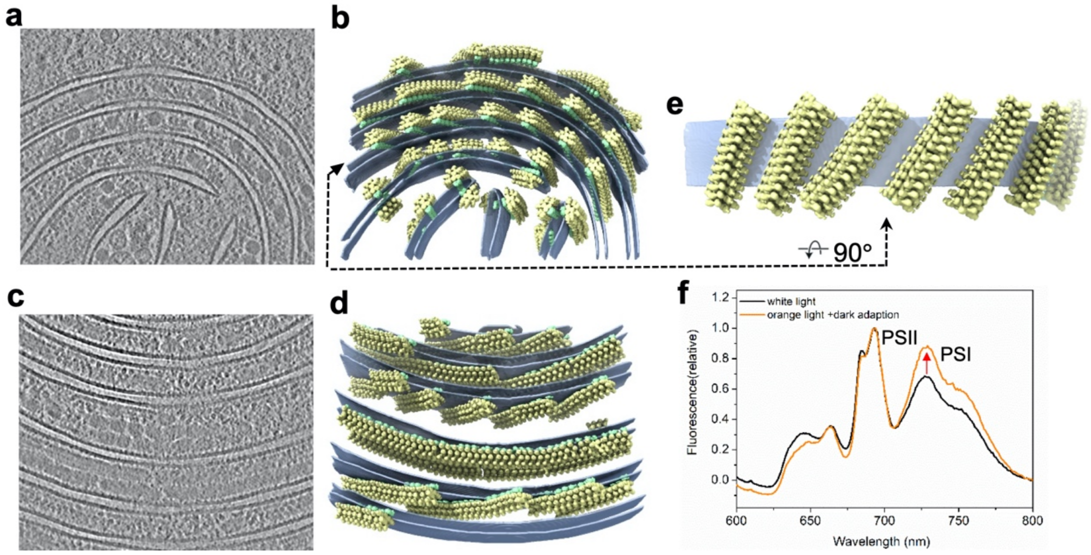


**Extended Data Figure 3 | The distribution of PBS-PSII supercomplexes in cell and 77 K fluorescence emission spectra of *S. platensis*.**

**a-d**, Two tomographic slices and the corresponding reconstructed tomograms (selected from 94 tomograms) with repositioned PBS-PSII supercomplexes showing the distribution of PBS-PSII in cell. PBSs and PSII dimers are colored in yellow and lightgreen, respectively. Thylakoid membrane is represented as transparent density layers. **e**, One of layers of (**d**) showing the top view of PBS-PSII supercomplexes in the membrane. **f**, Low temperature (77 K) fluorescence emission spectra of *S. platensis* cells cultured under low white light and orange light plus dark adaptation. The excitation wavelength was 580 nm.

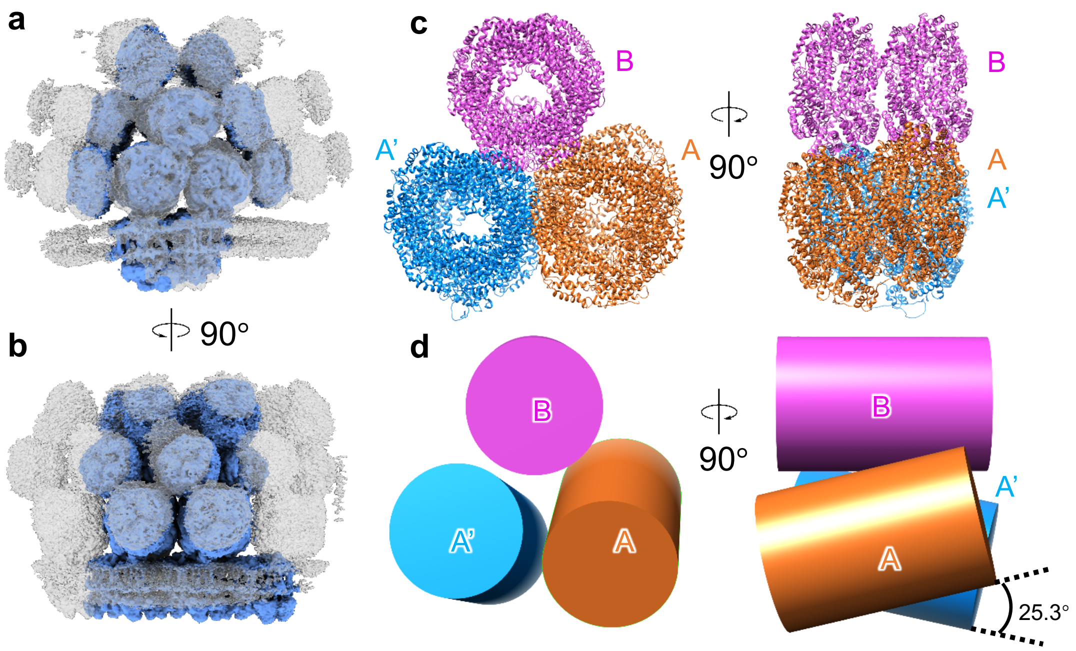


**Extended Data Figure 4 | Low-resolution cryo-EM density map of PBS-PSII supercomplex and PBS core structure.**

**a, b**, Different views of a low-resolution cryo-EM density map (5.1 Å) of PBS-PSII supercomplex showing the flexible rods. The stable region corresponding to the overall PBS-PSII model is colored in cornflower blue. **c**, Structure of PBS core without linker proteins. Core B, A and A' are colored in magentas, blue and orange, respectively. **d**, Schematic showing the slight twist (~ 25.3°) between two basal cylinders (A and A').

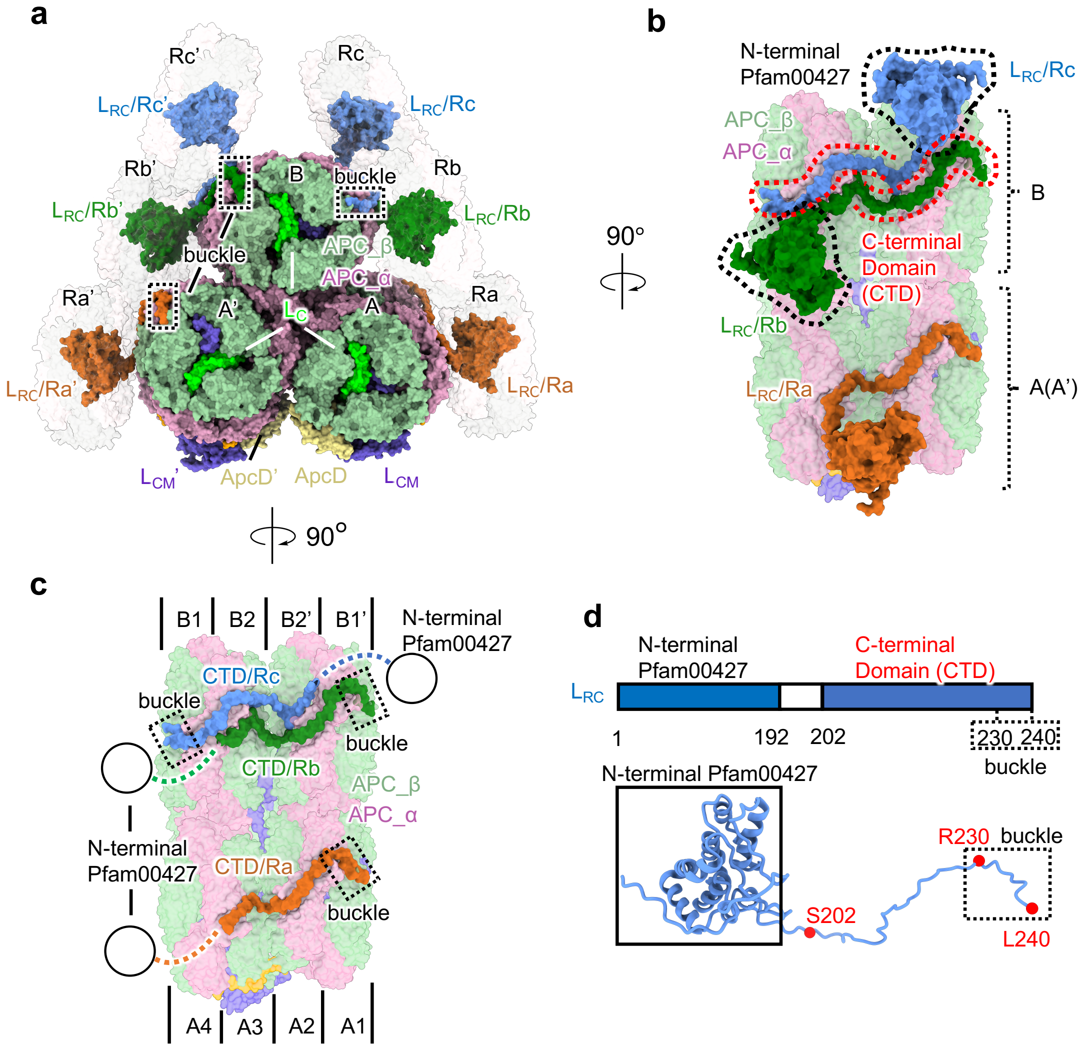


**Extended Data Figure 5 | Overall structure of *in situ* cyanobacterial PBS.**

**a-c**, Overall structure of *in situ* cyanobacterial PBS. The rods are shown as surface representation in 80% transparency. $L_{RC}$ of Ra/Ra', Rb/Rb' and Rc/Rc' are colored in orange, forest and cornflower blue, respectively. Core subunits are the same colors as those in Fig. 1a. **d**, The diagram of structural elements of $L_{RC}$ (upper panel) is shown above the structure (lower panel).

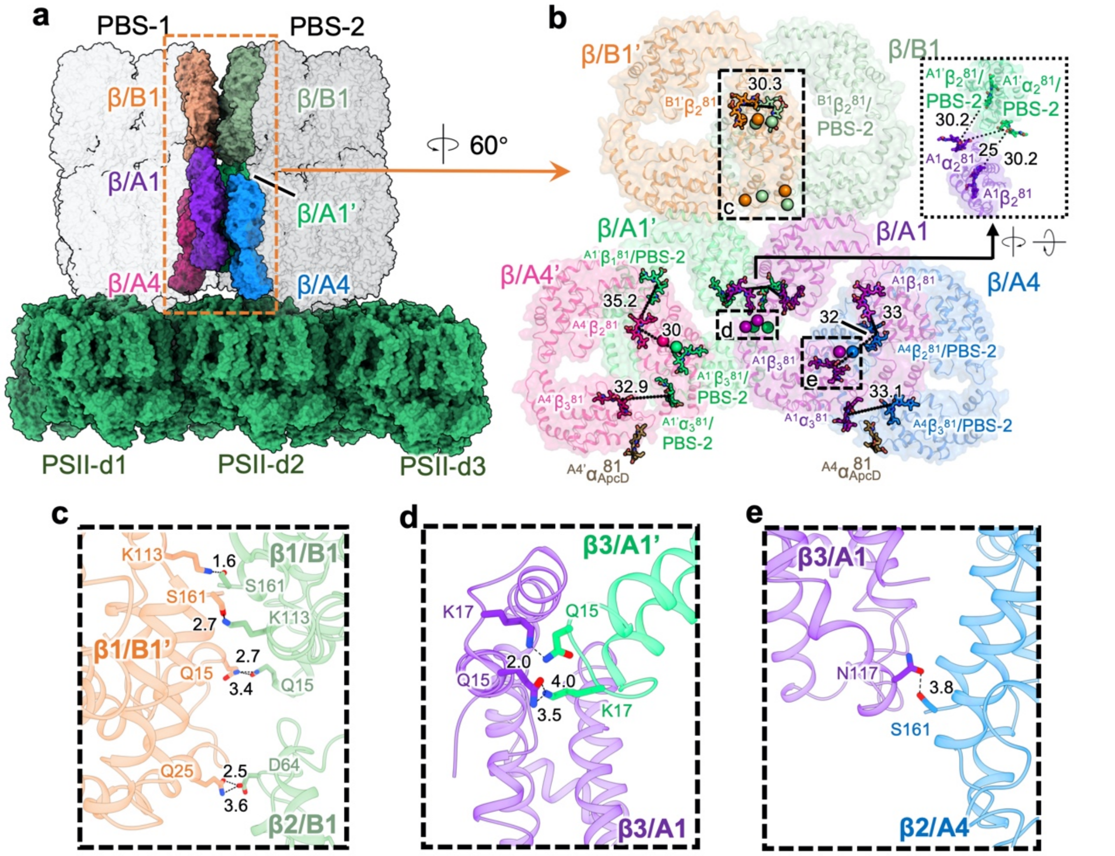


**Extended Data Figure 6 | Assembly of super-PBS and key billins mediating energy transfer in super-PBS.**

**a**, Overview of arrangement of super-PBS on the PSII dimers array. The trimeric β layers at the interface between PBS-1 and -2 cores are highlighted in different colors. **b**, Magnified view of the interfaces with the key bilins in different β layers. Key residues involved in the interface, rotated 60° relative to **a,** with key residues shown as spheres and key bilins as sticks. The numbers indicate the distances (Å) between the bilin pairs. **c-e**, The enlarged views show the details of the interactions between B1'/PBS-1 and B1/PBS-2 layers (**c**), A1/PBS-1 and A1' PBS-2 layers (**d**), and A1 PBS-1 and A4 PBS-2 layers (**e**).

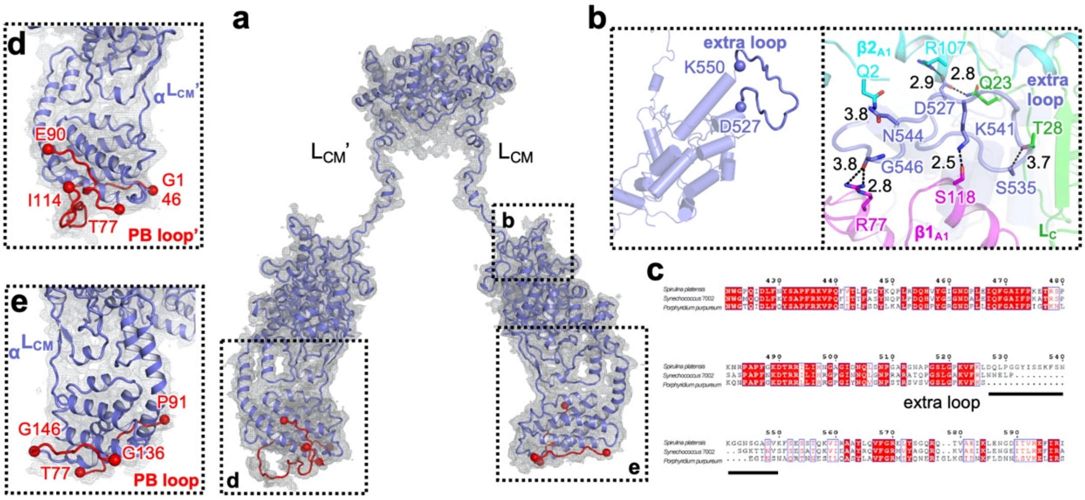


**Extended Data Figure 7 | Structure of the $L_{CM}$ and $L_{CM}$' from *S. platensis*.**

**a**, The density (mesh) for the linker proteins $L_{CM}$ and $L_{CM}$' superimposed with their atomic models (cartoon). **b**, The extra loop of $L_{CM}$ – D527 to K550 – shows several H-bonds with A1 layer and $L_C$. **c**, Sequence alignment of $L_{CM}$ from cyanobacteria and red algae shows cyanobacterial $L_{CM}$ has been evolved an extra loop region (D527 to K550). **d, e**, Close-up views of the PB loop of $L_{CM}$ (**d**) and $L_{CM}$' (**e**).

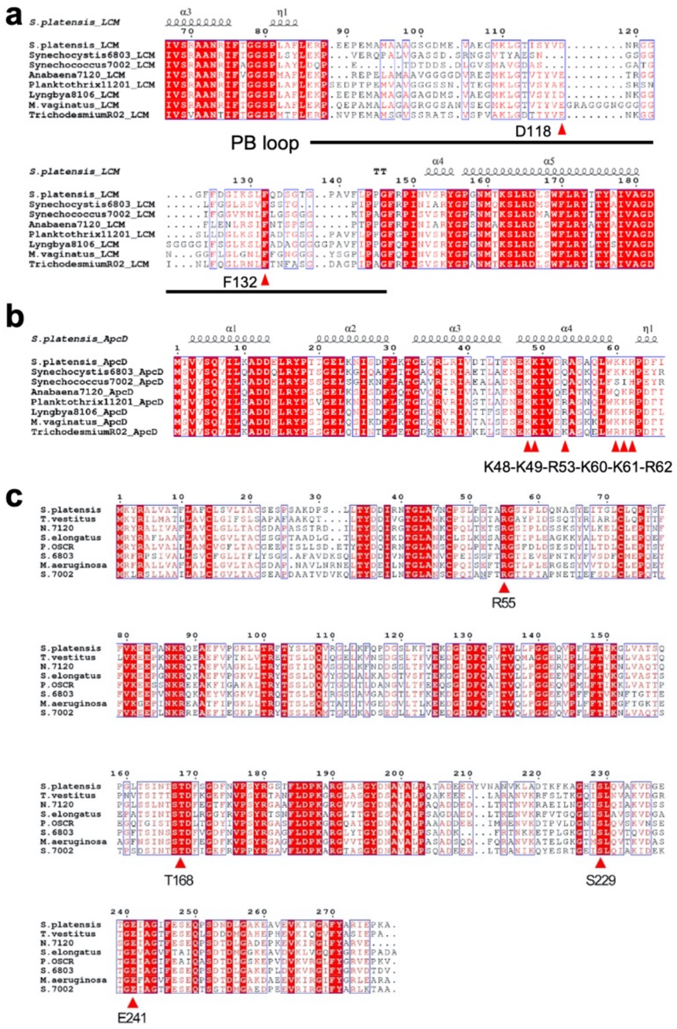


**Extended Data Figure 8 | Sequence alignment of $L_{CM}$, ApcD and PsbO from different cyanobacteria.**

**a-c**, Sequence alignments of $L_{CM}$ (**a**), ApcD (**b**) and PsbO (**c**) from *S. platensis* and other cyanobacteria. Used species are *S. platensis*, *Synechocystis* sp. PCC 6803, *Synechococcus* sp. Strain PCC 7002, Anabaena 7120, *Planktothrix* strain PCC 11201, *Lyngbya* aestuarii PCC 8106, *Microcoleus* vaginatus, *Trichodesmium* R02, *Taxillus vestitus*, *Synechococcus elongatus*, *Phormidium* sp. OSCR and *Microcystis aeruginosa*.

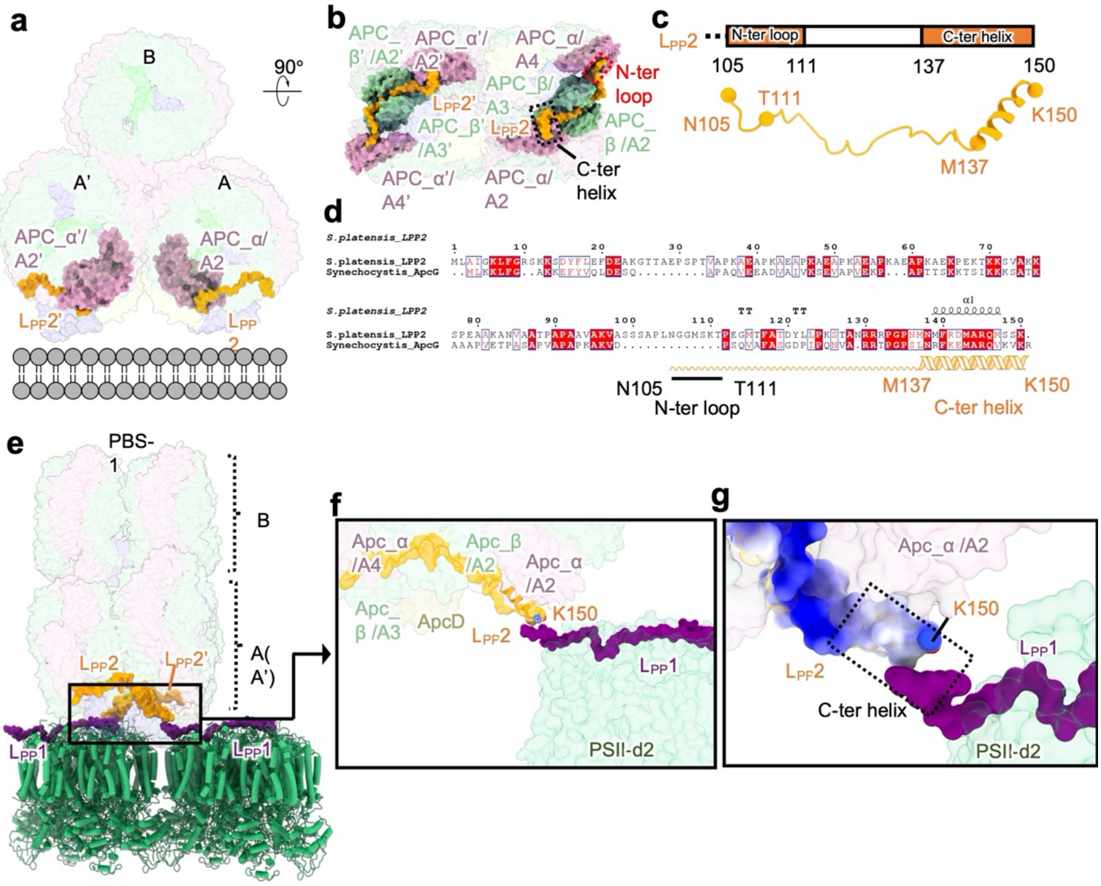


**Extended Data Figure 9 | Characterization of $L_{PP}2$.**

**a, b**, Two views showing the positions of $L_{PP}2$ and $L_{PP}2$' within the PBS. **c**, The diagram of structural elements of $L_{PP}2$ (upper panel) is shown above the structure (lower panel). **d**, Sequence alignment of $L_{PP}2$ with the homolog ApcG from *Synechocystis* sp. PCC 6803. **e-g**, Interactions between $L_{PP}2$ and $L_{PP}1$. $L_{PP}2$ in (**g**) is shown as electrostatic surface representation.

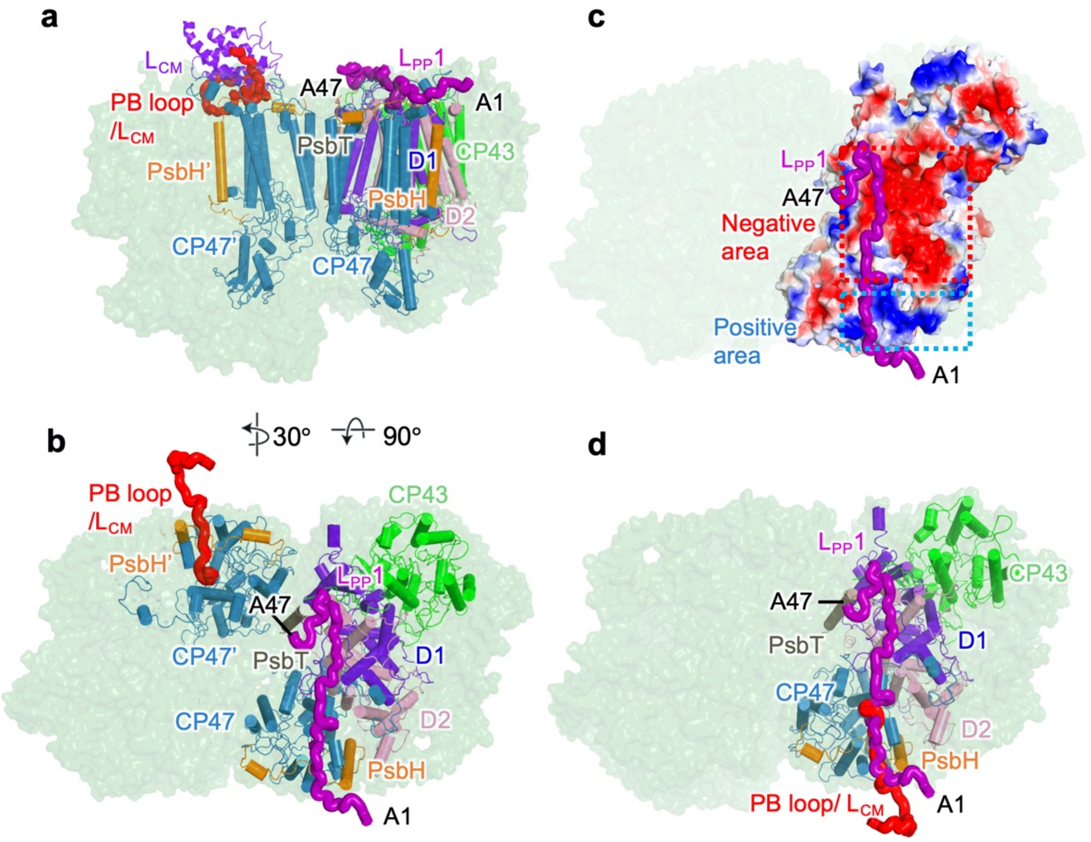


**Extended Data Figure 10 | Position of $L_{PP}1$ within the PSII dimer.**

**a, b**, Different views showing the position of $L_{PP}1$. **c**, Electrostatic interaction of $L_{PP}1$ with PSII. **d**, Superposition of the PSII-binding motif of PB loop/$L_{CM}$ and $L_{PP}1$. PB loop/$L_{CM}$ and $L_{PP}1$ are shown as surface representation and colored in red and purple, respectively.

### Extended Data Table 1. Cryo-EM data collection, refinement and validation statistics

| | PBS-PSII (EMDB-34487) (PDB 8H4W) | PBS-PSII (EMDB-34521) | PBS (EMDB-33605) | PSII dimer (EMDB-33597) |
|---|---|---|---|---|
| **Data collection and processing** | | | | |
| Magnification | | 53,000 | 53,000 | 53,000 |
| Voltage (kV) | | 300 | 300 | 300 |
| Electron exposure (e-/Å$^2$) | | 35 | 35 | 35 |
| Defocus range (μm) | | -1.0 ~ -6.0 | -1.0 ~ -6.0 | -1.0 ~ -6.0 |
| Pixel size (Å) | | 1.632 | 1.632 | 1.632 |
| Symmetry imposed | | C1 | C1 | C1 |
| Initial particle image (no.) | | 167,000 | 1,873,000 | 664,000 |
| Final particle image (no.) | | 127,000 | 168,000 | 83,000 |
| Map resolution (Å) | | 5.1 | 3.6 | 3.5 |
| FSC threshold | | 0.143 | 0.143 | 0.143 |
| Map resolution range (Å) | | 5.1 ~ 20 | 3.6 ~ 8.0 | 3.5 ~ 7.8 |
| **Refinement** | | | | |
| Initial model used (PDB code) | 7EXT, 7RCV | | | |
| Model resolution (Å) | 3.6 | | | |
| FSC threshold | 0.143 | | | |
| Model resolution range (Å) | 3.5 ~ 11 | | | |
| Map sharpen *B* factor (Å$^2$) | | -136 | -99 | -90 |
| Model composition | | | | |
| Non-hydrogen atoms | 1,264,870 | | | |
| Protein residues | 153,311 | | | |
| Ligands | 2,436 | | | |
| *B* factors (Å$^2$) | | | | |
| Protein | 89.29 | | | |
| Ligands | 98.55 | | | |
| R.m.s.deviations | | | | |
| Bond lengths (Å$^2$) | 0.011 | | | |
| Bond angles (°) | 2.058 | | | |
| Validation | | | | |
| MolProbity score | 2.1 | | | |
| Clashscore | 11.54 | | | |
| Poor rotamers (%) | 1.45 | | | |
| Ramachandran plot | | | | |
| Favored (%) | 94.06 | | | |
| Allowed (%) | 5.77 | | | |
| Disallowed (%) | 0.17 | | | |
